# VASAL: A Vascular Substrate Algorithm to Generate Microvascular Network Phantoms

**DOI:** 10.1101/2025.09.05.674056

**Authors:** Elizabeth Powell, Geoff J. M. Parker, Marco Palombo

## Abstract

To date, few methods have been proposed for synthesising realistic microvascular networks, particularly with compatibility for diffusion MRI (dMRI) simulations. This work presents a generative algorithm for building computational brain microvascular network phantoms, together with a Monte Carlo framework for simulating spin flow dynamics and corresponding dMRI signals. Phantom morphology can be user-tuned according to the: number of generations (branches); total number of segments; segment diameter and length; angle between connecting segments, and; maximum extent in space.

The geometric fidelity of resulting VASAL phantoms was validated against an exemplar rat hippocampal network. Comparable distributions of segment lengths, diameters and orientations, and branching characteristics were obtained, demonstrating VASAL replicated realistic microvascular structures. Implementation of spin flow dynamics was validated by simulating dMRI signals in the ballistic regime, which has a well-defined analytical solution; agreement was excellent between simulated signals and analytical solutions (average root mean square error of 0.005).

Following this validation of network geometry and spin flow dynamics, VASAL phantoms with different morphologies were generated to evaluate key IVIM model assumptions. Phantom geometry substantially affected dMRI signals and caused deviations from the IVIM model: for example, complex branching phantoms displayed non-Gaussian signal attenuation and diffusion time dependence. While the IVIM model was exemplified here, the presented VASAL algorithm holds potential for advancing other dMRI acquisitions and models: by enabling simulation of complex blood flow within physiologically accurate vascular networks, a route to disentangling how vascular structure and function influence dMRI signals is initiated.

## I. Introduction

Microstructural imaging techniques aim to non-invasively infer microscopic tissue features from macroscopic images, ultimately reducing the need for invasive biopsies and histological procedures. Diffusion MRI (dMRI) - currently the dominant non-invasive microstructural imaging technique - can be used to infer tissue properties such as the diameter and orientation of neuronal axons, the size and density of cells, or the permeability of cell membranes [1]. The diffusive motion of water molecules (spins) is influenced by such features, and can be probed in dMRI by the application of magnetic field gradients, typically along multiple directions. However, the complexity and heterogeneity of biological tissues means that the relationship between acquired dMRI signals and underlying tissue features is far from trivial. Numerical phantoms and simulations have emerged as a powerful tool for disentangling the influences of different tissue features on MRI signals, but the focus to date has primarily been on white [2]–[6] and grey [7], [8] matter models; few in silico capillary network phantoms exist, despite the importance of the brain microvasculature.

Vascular alterations are implicated in numerous neurological conditions, including brain tumours [9], stroke [10], cerebral small vessel disease [11], and dementia [12]. The dMRI signal can be sensitised to blood via the pseudo-diffusivity effect, which describes blood flow through tortuous vessels as a random walk akin to true diffusion processes but on a different spatio-temporal scale. For example, the IV IM (lntraVoxel Incoherent Motion) [13] technique has been used to assess tissue perfusion following stroke [14], VERDICT-MRI (Vascular, Extracellular, and Restricted Diffusion for Cytometry in Tumours) [15] has been adapted for brain tumour characterisation [16], and double diffusion encoding methods have been developed for imaging blood-brain barrier permeability [17], [18].

Each of these techniques relies on a constrained biophysical model to reconstruct tissue feature maps, imposing assumptions and simplifications that are still under debate. Validation of these methods and their assumptions using ex vivo imaging (histology) or in vitro models can be valuable, but such methods are impractical for replicating the wide range of pathophysiological conditions seen across disease and often suffer from localised sampling that can be challenging to relate to the large volumes sampled using dMRI. Physical phantoms [19] can provide controllability and can be produced at a scale matching dMRI sampling volumes, but are time consuming, expensive to develop, and make use of simplifying assumptions themselves, therefore limiting their value.

Imaging-derived numerical phantoms [20]–[23], although lacking a generative component for controlling vascular network morphology, are useful for deriving target geometric characteristics. Previous studies in this area reveal the following principal features of tissue vascular microstructure:

1. Microvascular networks (consisting of arterioles, venules and capillaries) form ‘mesh-like’ structures that are homogenous and space-filling over a characteristic length scale (25-75 µm [24]), with approximately convex extravascular domains [25];
2. Arterio-venous networks are quasi-fractal, composed of branching ‘tree-like’ structures [24];
3. Vascular networks of any scale have vertices with mainly three connections [26];
4. Microvessel diameters range between 4-20 µm.

Few generative computational methods have been proposed for synthesising microvascular networks, especially with compatibility for dMRI simulations. One used long, randomly folded tubes [27], but did not allow for more complex geometries such as branching. Methods using Voronoi diagrams to create lattice-like meshes [25] or diffusion-limited aggregation [28] have explicitly modelled branching, but imposed uniform capillary diameters. A common alternative for simulating micovascular MR signals without explicitly generating numerical phantoms is to model capillary segments as disconnected, randomly-oriented, straight cylinders [29], [30], but this also does not allow for complex geometries. Recently, an algorithm synthesising large-scale vascular networks was proposed [31], but has limited flexibility to modulate microvessel features.

This work presents a generative algorithm for computational brain microvascular phantoms with flexible network morphology, and models blood flow effects on dMRI signals. The paper is organised as follows: Sec. II summarises background theory on flow and the dMRI signal; Sec. III introduces VASAL (the VAscular Substrate ALgorithm), developed here for generating brain microvascular phantoms; Sec. IV describes the implementation of flow dynamics through VASAL phantoms; Sec. V validates achieved phantom morphologies and the implementation of flow dynamics; Sec. VI evaluates phantom morphology effects on IVIM signals.

## II. MRI Theory

Diffusion MRI conventionally utilises the pulsed gradient spin echo (PGSE) sequence to encode local water diffusion properties. To recap the theory behind PGSE signal generation: spin situated in a magnetic field gradient ***g*** (*t*) acquires a spatially-dependent phase

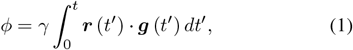

where ***r*** (*t*^*′*^) is the time-dependent spin position and *γ* is the gyromagnetic ratio. A distribution of phase shifts, *P* (*ϕ, t*), in a system of spins attenuates the MR signal according to

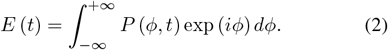

For spins undergoing unrestricted diffusion, the phase distribution function is typically Gaussian:

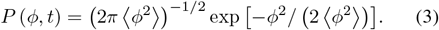

Using a standard integral (3.323(2) in [32]), the signal attenuation for a Gaussian phase distribution becomes

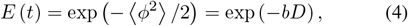

where the phase distribution variance for a PGSE is [33]

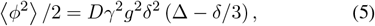

with the *b*-value given by the gradient pulse duration, *δ*, and separation, Δ:

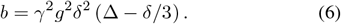

For spins undergoing translational motion owing to flow, Eq. 1 can be written in terms of spin velocity, ***v***, and time, *t*:

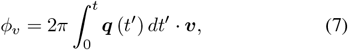

where the dephasing magnitude of the gradient is given by ***q*** = (2*π*)^*−*1^ *γδ****g***. For plug flow along the gradient direction, all spins accrue the same phase shift and there is no signal attenuation. For a distribution of flow velocities, *P* (*v*), the signal attenuation is described by

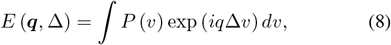

### Ballistic regime

For a uniform velocity distribution, Eq. 8 reduces to

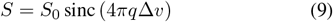

A uniform velocity profile can arise for plug flow in a network of randomly-oriented capillary segments if spins do not change direction during the diffusion time, i.e. for slow flow, long segments, or short diffusion times; these conditions define the ballistic regime. A diffraction pattern can thus be observed in the signal, with minima occurring when spin displacement correlates with the segment length scale. Functionally, minima occur when

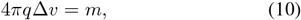

where *m* ∈ ℤ_≠0_. Deviations from a uniform velocity distribution, arising, for example, from laminar flow or gradient pulses with finite length, blur this diffraction pattern.

### Diffusive regime

In this regime, where blood water spins change direction multiple times during the diffusion time (i.e. for fast flow, short capillary segments, or long diffusion times), the signal attenuation from flow becomes mono-exponential:

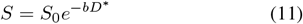

where the pseudo-diffusion coefficient, *D*^*∗*^, is related to the segment length scale, *l*, and mean spin velocity, ⟨*v*⟩ [13]:

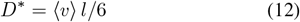

## III. VASAL: Vascular Substrate Algorithm

VASAL has three main steps: (i) defining capillary segment morphology given a set of user-input characteristics; (ii) growing the vascular tree, and; (iii) generating a 3D mesh to be used in Monte Carlo simulations. The algorithm is implemented in Matlab (The Mathworks, R2024b).

### A. Defining Morphological Characteristics

Phantom morphology (i.e. the simulated capillary bed characteristics) is tuned according to the following user-input variables: (i) the number of generations, *g*; (ii) target total number of segments, *N*; (iii) minimum segment diameter, *d*_*min*_; (iv) minimum mean segment length, 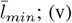 minimum angle between connecting segments, *θ*_*min*_, and; (vi) maximum extent in space, L = [X,Y,Z] (i.e. ‘voxel’ size). The minimum velocity, *v*_*min*_, is also user defined.

To maintain flow conservation and geometric self-similarity across branch generations, the following constraints - from Henkelman et al. [29] - are enforced. Vessel diameters, which change only at bifurcations, follow the relationship [29]:

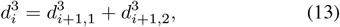

where *i* indexes the vessel generation, *d*_*i*_ is the parent vessel diameter and *d*_*i*+1,1_, *d*_*i*+1,2_ are the child diameters. Without loss of generality, both child vessels are assigned the same diameter here (*d*_*i*+1,1_ = *d*_*i*+1,2_). Segments are assigned a random length taken from a normal distribution around the mean value, 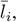 which, to preserve geometric self-similarity at each branch generation, scales according to vessel diameter as 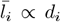 [29]. It follows that the total blood volume in each generation must also be a constant, leading to the condition that the number of vessel segments in each generation, *N*_*i*_, scales with diameter as 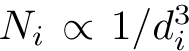 [29]. Finally, a constant flow *F* is enforced across branch generations [29]:

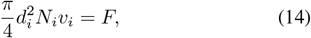

where *v*_*i*_ is the blood velocity in the *i*th generation.

### B. Growing the Vascular Tree

Vascular networks can be designed either with periodic boundary conditions (i.e. with inlets/outlets) or with effectively reflective boundary conditions (closed loop, no inlets/outlets).

The number of unique network paths is defined by the number of branch generations, *g*: *N*_*path*_ = 2^*g−*1^ (Fig. 1). Each path is grown sequentially, retaining segments from previous generations and forging new ones from the most recent branch node. Vessel growth is a non-Markovian stochastic process resulting in a self-avoiding walk (SAW); vessel growth may therefore be terminated if no new segments can be generated without intersecting existing segments. However, the SAW condition can be relaxed to produce true random-walk networks, if needed. Vessels are grown as follows (Fig. 1):

**Fig. 1.**
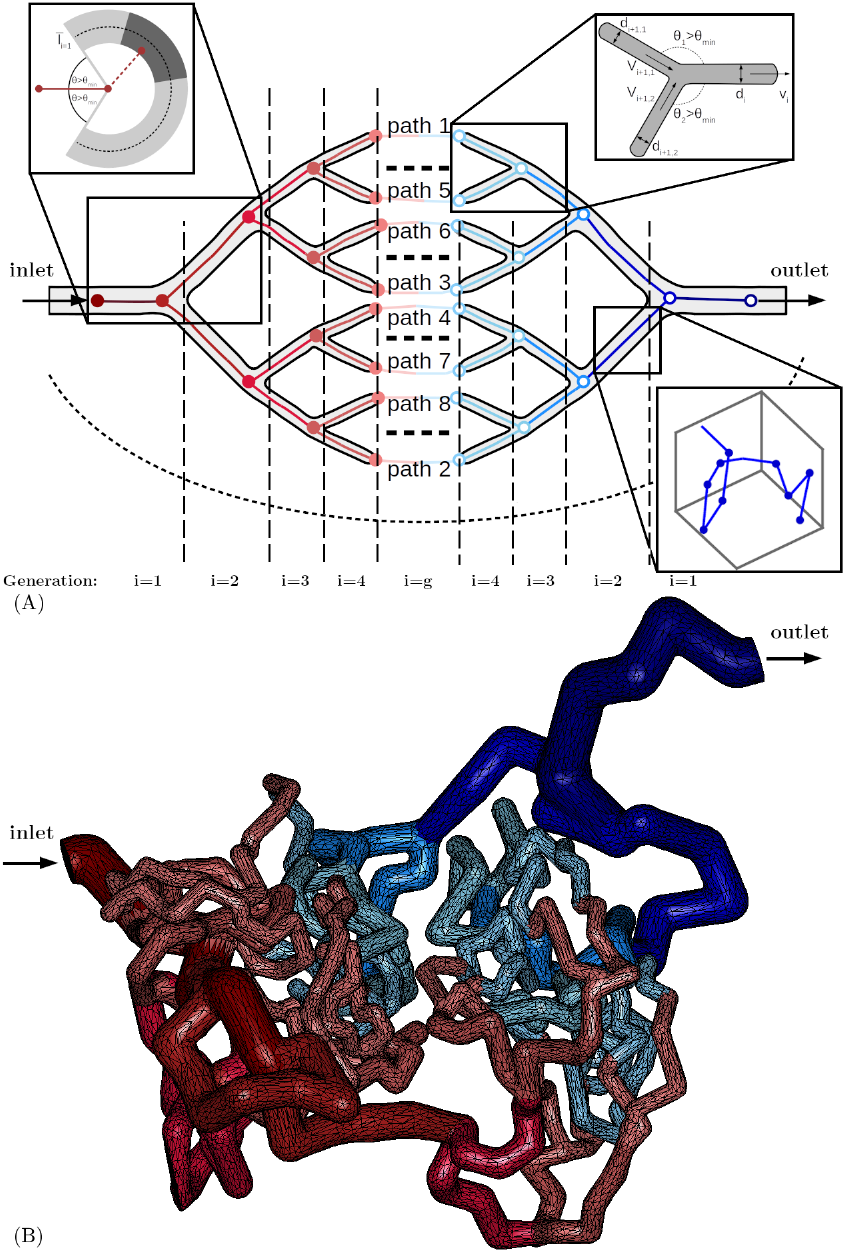
VASAL schematic. **(A)**. Paths are generated sequentially; for example, path 1 is grown in its entirety from inlet to outlet, path 2 uses the existing skeleton at ***i* = 1** and new segment generation begins at ***i* = 2**, and so on. All generations contain multiple segments (inset, bottom right). Branch nodes are represented by closed circles, terminal branch nodes by open circles. Segments are generated from a prospective node within the region satisfying segment length and connecting angle statistics; this region is highlighted in light grey (inset, top left), while the dark grey region shows how connecting angle statistics can be modified to guide segment growth towards the terminal node. Each segment has an associated velocity vector oriented along its axis (inset, top right). **(B)**. An exemplar VASAL mesh.

1. Inlets/outlets are placed at the same (user-defined) location on opposing voxel faces (periodic boundary conditions) or otherwise at the same location within the voxel (reflective boundary conditions);
2. Prospective nodes (25 initially) are randomly generated according to segment length and connecting angle statistics 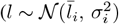 and *θ > θ*_*min*_, respectively).
3. A segment is added to the skeleton if it connects to a proposed node without intersecting existing segments.
4. If all prospective nodes intersect with the existing skeleton, a new set are generated; this step is repeated up to ten times, doubling the number of generated nodes each time (balancing efficient spatial exploration while avoiding infinite node placement searches).
5. If no new node is accepted, the current node is discarded and Steps 1–4 are repeated from the previous node.
6. Branch nodes are defined when the path segment count is 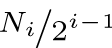. Path growth continues along one branch through all generations (recording subsequent branch nodes) until the terminal node is reached. Segments for the next path are grown from its latest branch node.
7. Steps 1-7 are iterated until all paths are complete.

To ensure closed networks, connecting angle statistics in Step 2 are modified if the current-to-terminal node distance exceeds the effective random walk length for the remaining segments; then, inertia toward the terminal node is enforced by restricting prospective nodes to a solid angle oriented at the terminus.

The vascular tree skeleton is saved as a SWC file, with each segment in the *i*th generation assigned a diameter *d*_*i*_ (Eq. 13).

## C. Rendering the 3D Mesh

Blender (https://www.blender.org/) and the SWC mesher addon (https://github.com/mcellteam/swc_mesher) are used to generate a triangulated surface mesh around the skeleton. Metaballs with minimum/maximum diameters matching the user-defined vessel diameters, mesh resolution = 0.1 × *d*_*min*_ and scale factor = 1.5385 are placed along the skeleton; values were set empirically to ensure neighbouring metaballs fuse and produce smooth surfaces with correct segment diameters. The mesh is downsampled to 5% of original triangles to reduce the face count and thus computation time in simulations; mesh fidelity was not impacted as assessed visually.

## IV. Flow Dynamics

A Monte Carlo framework tailored for simulating dMRI signals was implemented in Matlab 2024b (The Mathworks). Spin diffusion dynamics and resulting phase accrual during a PGSE experiment were modelled based on Hall & Alexander’s [34] framework; this was extended here to model flow concurrently.

First, a velocity field was created for the VASAL phantom. Each segment was assigned a velocity vector oriented along its axis (directed from inlet to outlet) with a magnitude governed by its diameter and user-input minimum velocity, *v*_*min*_ (Eq. 14). All mesh faces of a segment were assigned its velocity vector, rotated if necessary to ensure parallel alignment with the face (i.e. at corners/bifurcations). This effectively models plug flow of non-interacting spins, which is a good approximation for capillary blood flow; however, more complex velocity fields could be calculated and imported instead.

An intravascular spin by the *j*th face acquires a flow drift:

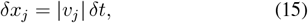

where *δt* is the Monte Carlo time step. Total spin displacement during *δt* is a combination of diffusion and flow:

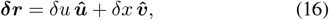

where ***û*** and ***v̂*** are unit vectors representing the direction of diffusion (random in space) and flow (along vessel segments) respectively, with *δu* the diffusion step size determined by the intrinsic diffusion coefficient *D* as 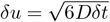 (in 3D), and *δx* = *δx*_*j*_ intravascular and *δx* = 0 everywhere else.

## V. Validation

This section outlines experiments: (i) validating phantom morphology, and; (ii) testing the new Monte Carlo flow simulation.

### A. Morphological Characteristics

#### 1) Methods

Micro-CT data obtained from a corrosion cast of a Wistar-Kyoto rat (aged 13 months) [22] were analysed to obtain representative microvasculature morphological characteristics. Briefly, the data comprised a binary segmentation of a 1.499 mm^3^ hippocampal section with 0.7499 µm^3^ resolution. An analysis framework was developed (described in Appendix A) to characterise geometric properties including segment lengths, diameters, and orientation distributions; these were used to inform and validate VASAL phantom morphology.

The analysis framework was also applied to ten VASAL phantoms (periodic, self-avoiding, *g* = 5, *N* = 350, *d*_*min*_ = 6 µm, *l*_*min*_ = 10 ± 4 µm, *θ*_*min*_ = 20^*°*^, *L* = [250, 250, 250] µm^3^; Table I) to ensure it returned a feasible network representation.

**TABLE I.** Effect of morphological characteristics on *D*^*∗*^. Phantom characteristics are summarised for each section, where ***g*** is the number of generations, ***N*** the number of segments, ***d***_***min***_ the minimum segment diameter, ***l***_***min***_ the minimum segment length set, 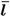 the mean segment length realised over all 10 meshes, ***θ***_***min***_ the minimum connecting angle, and ***L*** the maximum extent in space. Simulation parameters are also provided, where ***v***_***min***_ is the minimum velocity (corresponding to the smallest diameter), 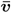 is the mean velocity across vessel diameters, ***D***^***∗***^ is the pseudo-diffusivity estimated from the signal attenuation, ***D***^***∗***^ (GT) is the theoretical pseudo-diffusivity calculated as ⟨***l***_**1:*N***_ ***v***_**1:*N***_ ⟩***/*6** (where ***l***_**1:*N***_, ***v***_**1:*N***_ are the lengths, velocities in all segments, respectively), and # seg. the mean number of segments traversed in the experiment time (**Δ + *δ***). The VASAL phantoms with morphology comparable to the rat hippocampus are highlighted in grey.

| Section | Network type | Morphological characteristics | | | | | | | Simulations | | | $D^*$ (GT)<br>[$\mu\text{m}^2/\text{ms}$] | # seg. |
| --- | --- | --- | --- | --- | --- | --- | --- | --- | --- | --- | --- | --- | --- |
| | | $g$ | $N$ | $d_{min}$<br>[$\mu\text{m}$] | $l_{min}$<br>[$\mu\text{m}$] | $\bar{l}$<br>[$\mu\text{m}$] | $\theta_{min}$<br>[$^\circ$] | $L$<br>[$\mu\text{m}^3$] | $v_{min}$<br>[mm/s] | $\bar{v}$<br>[mm/s] | $D^*$<br>[$\mu\text{m}^2/\text{ms}$] | | |
| V-A | periodic, SAW | 5 | 350 | 6 | $10 \pm 4$ | 14 | 20 | 250,250,250 | 2 | 3 | $13.9 \pm 1.3$ | 6.1 | 9.0 |
| V-B | periodic, RW | 1 | 350 | 6 | $10 \pm 4$ | 11 | 0 | 250,250,250 | 2 | 2 | $4.0 \pm 0.3$ | 3.7 | 8.9 |
| VI-A | periodic, RW | 1 | 350 | 6 | $5 \pm 2$ | 7 | 0 | 250,250,250 | 2 | 2 | $2.2 \pm 0.3$ | 2.4 | 12.7 |
| | periodic, RW | 1 | 350 | 6 | $10 \pm 4$ | 11 | 0 | 250,250,250 | 2 | 2 | $4.0 \pm 0.3$ | 3.7 | 8.9 |
| | periodic, RW | 1 | 350 | 6 | $15 \pm 6$ | 16 | 0 | 250,250,250 | 2 | 2 | $5.3 \pm 0.5$ | 5.2 | 6.6 |
| | periodic, RW | 1 | 350 | 6 | $20 \pm 8$ | 21 | 0 | 250,250,250 | 2 | 2 | $7.1 \pm 0.5$ | 6.9 | 5.2 |
| | continuous, RW | 1 | 50 | 6 | $10 \pm 4$ | 11 | 0 | 62,55,60 | 2 | 2 | $4.3 \pm 1.2$ | 3.7 | 8.8 |
| | continuous, RW | 1 | 100 | 6 | $10 \pm 4$ | 11 | 0 | 82,74,94 | 2 | 2 | $3.9 \pm 0.9$ | 3.7 | 8.9 |
| | continuous, RW | 1 | 350 | 6 | $10 \pm 4$ | 11 | 0 | 207,181,165 | 2 | 2 | $4.2 \pm 0.5$ | 3.7 | 8.7 |
| VI-B | periodic, RW | 1 | 350 | 6 | $10 \pm 4$ | 11 | 20 | 250,250,250 | 2 | 2 | $6.4 \pm 0.6$ | 3.7 | 8.8 |
| | periodic, SAW | 1 | 350 | 6 | $10 \pm 4$ | 11 | 0 | 250,250,250 | 2 | 2 | $10.3 \pm 0.6$ | 3.7 | 8.8 |
| | periodic, SAW | 1 | 350 | 6 | $10 \pm 4$ | 11 | 20 | 250,250,250 | 2 | 2 | $8.8 \pm 0.6$ | 3.7 | 8.8 |
| | periodic, RW | 5 | 350 | 6 | $10 \pm 4$ | 14 | 0 | 250,250,250 | 2 | 3 | $7.6 \pm 1.3$ | 6.3 | 9.0 |
| | periodic, SAW | 5 | 350 | 6 | $10 \pm 4$ | 14 | 20 | 250,250,250 | 2 | 3 | $13.9 \pm 1.3$ | 6.1 | 9.0 |
| VI-C | periodic, SAW | 5 | 350 | 6 | $10 \pm 4$ | 14 | 20 | 250,250,250 | 2 | 3 | $13.9 \pm 1.3$ | 6.1 | 9.0 |

#### 2) Results

Fig. 2 shows the morphological characteristics of the VASAL phantom (ground truth and estimated using the framework in Appendix A) and those estimated in the rat data. Good agreement was observed between the ground truth and estimated VASAL phantom morphology, indicating the analysis framework returned realistic parameters (Fig. 2A). Good agreement was also seen between most features of the VASAL and rat meshes (Fig. 2B); however, there were differences in the estimated segment diameters. As the rat network contained some larger vessels as well as the capillary bed, this may explain the higher incidence of larger diameters in the rat data.

**Fig. 2.**
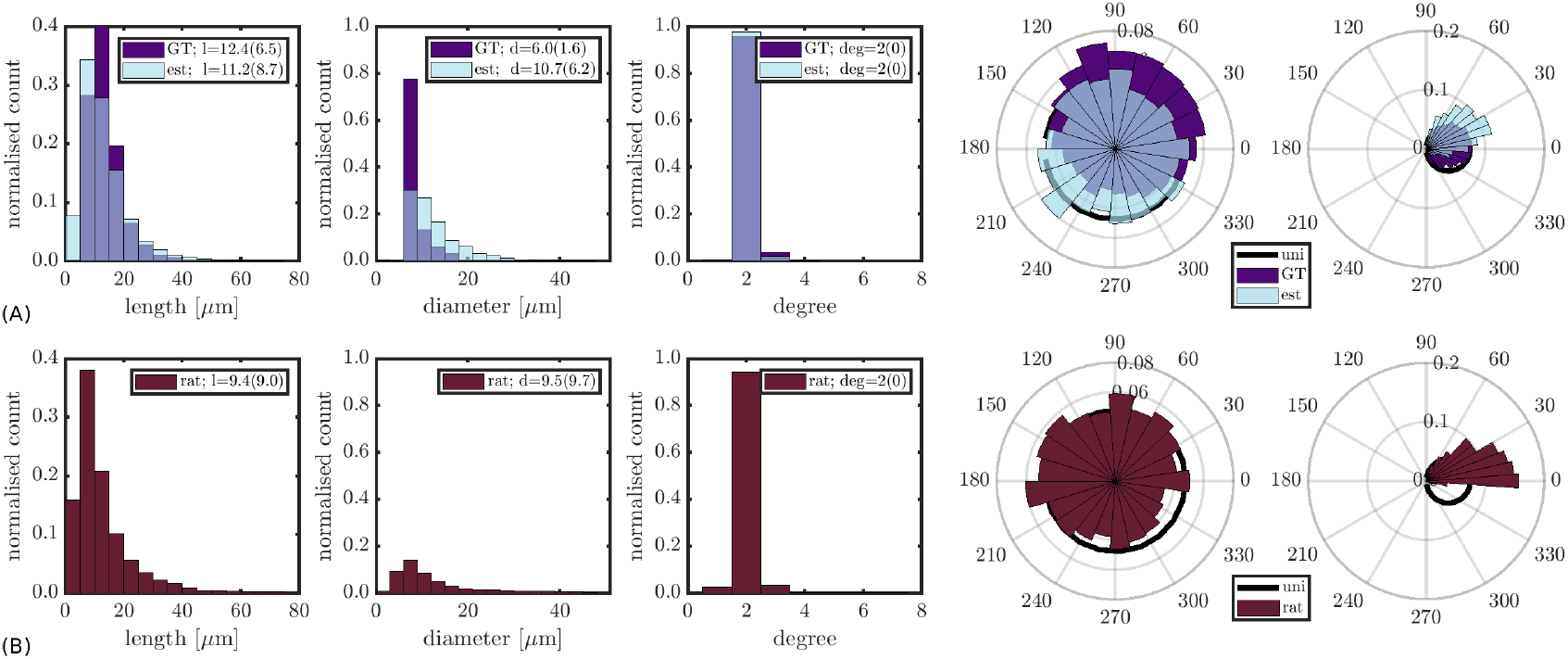
Validating VASAL phantom morphological characteristics. **(A)**. Distributions of segment length, diameter, connection degree and orientation are shown for the VASAL phantoms, with ground truth characteristics shown in purple and estimated characteristics in blue. **(B)**. Distributions of segment length, diameter, connection degree and orientation estimated from the rat brain microvasculature.

### B. Flow Dynamics

#### 1) Methods

Flow only (no diffusion) was simulated in 1000 infinite cylinders uniformly oriented on the sphere (length ≫ flow distance) (Fig. 3A) and in 10 periodic, non-branching, self-intersecting VASAL meshes (i.e. resembling random walks) generated with *g* = 1, *N* = 350, *d* = 6 µm, *l* = 10 ± 4 µm, *θ*_*min*_ = 0^*°*^, *L* = [250, 250, 250] µm^3^ (Table I, Fig. 3B). While these are two very different systems, identical dMRI signals should be obtained in the ballistic regime; this setup can therefore validate both the implementation of flow dynamics (as infinite random cylinders have a well-defined analytical solution) and the fidelity of VASAL phantom morphology under known conditions.

**Fig. 3.**
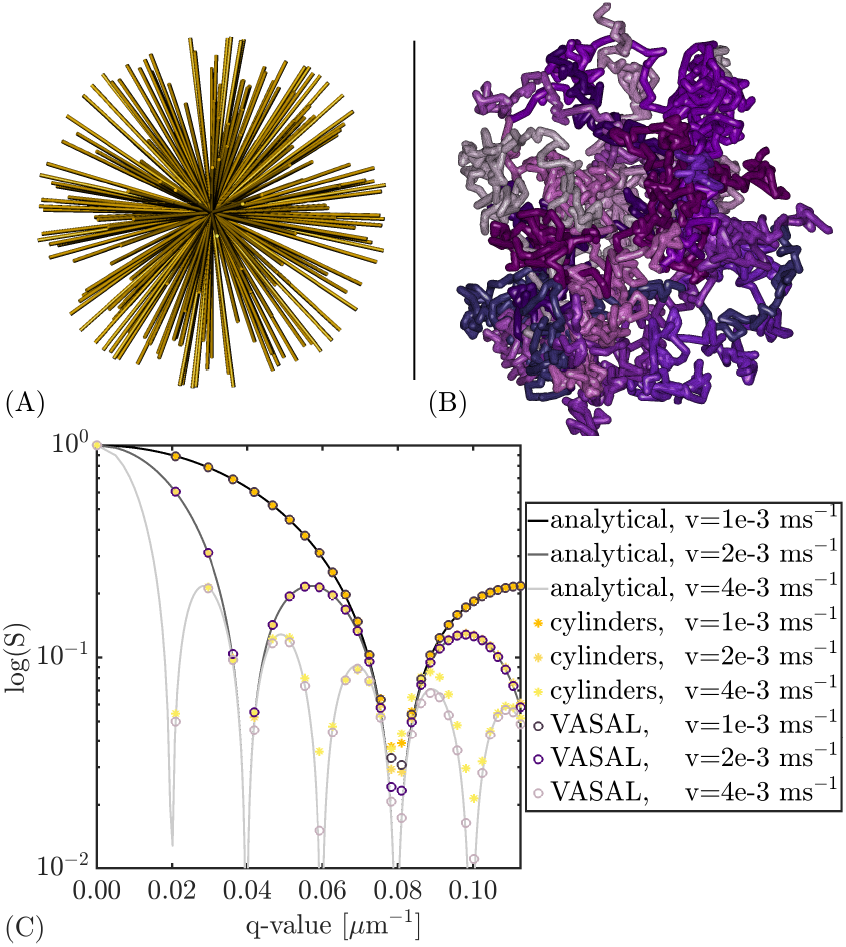
Validating flow dynamics. **(A)**. A subset of the 1000 cylinder meshes. **(B)**. All 10 VASAL phantoms, each in a different shade. **(C)**. Signal-vs-***q*** plots are shown for each flow velocity for the analytical solution (solid lines), and for spin trajectories in the cylinders (yellow markers) and VASAL phantoms (purple markers).

For the cylinders, 50 spins per cylinder (50,000 total) were initialised uniformly in a plane at their centres, preventing spins reaching the ends; for the VASAL phantoms, 5,000 spins per phantom (50,000 total) were initialised uniformly inside. Spin trajectories were generated with time step interval *δt* = 1 ms (for computational efficiency in the absence of diffusion, this is the maximum *δt* possible given the gradient separation, below) and three flow velocities *v* = [1, 2, 4] mm/s. No spin dynamics were simulated outside the meshes.

Signals were computed using the short gradient pulse approximation (SGP) (i.e. *δ* ∼ 0), with Δ = 1 ms (ensuring the majority of spins remained in their initial VASAL segment), 30 *b*-values uniformly spaced between *b* = 0 − 0.5 ms/µm^2^ (equivalently *q* = 0 − 0.1125 µm^*−*1^), and 32 gradient directions per *b*-value. Signals were averaged over gradient directions for all 1000 cylinders and all 10 VASAL phantoms.

#### 2) Results

Fig. 3C shows the signals simulated in the cylinder and VASAL phantoms. Both show excellent agreement with the analytical solution (Eq. 9): root mean squared errors (RMSE) for the three velocities were RMSE_*cyl*_ = [0.006, 0.005, 0.012], and RMSE_*vasal*_ = [0.005, 0.005, 0.004] for the cylinder and VASAL meshes respectively.

## VI. IVIM Experiments

This section utilises VASAL and the new Monte Carlo flow simulation framework to explore the effect of capillary network morphology on IVIM signals, one of the most popular dMRI methods to quantify blood perfusion in vivo.

### A. True Random Walks

#### 1) Methods

A key IVIM model assumption is that blood flow in capillaries mimics the random walk of diffusion processes. To best mimic this condition, periodic, non-branching, self-intersecting VASAL phantoms were generated with *g* = 1, *N* = 350, *d* = 6 µm, *θ*_*min*_ = 0^*°*^, *L* = [250, 250, 250] µm^3^, and *l*_1_ = 5 ±2 µm, *l*_2_ = 10 ±4 µm, *l*_3_ = 15 ±6 µm, *l*_4_ = 20 ±8 µm (10 phantoms per segment length, *l*) (Table I).

The number of segments required to adequately approximate the statistical characteristics of a random walk was also investigated. Continuous (ensuring no restrictive effects from pre-defining the phantom extent in space, *L*), non-branching, self-intersecting VASAL phantoms were generated with *g* = 1, *d* = 6 µm, *l* = 10 ±4 µm, *θ*_*min*_ = 0^*°*^, and *N* = [50, 150, 250, 350] (10 phantoms per segment number, *N*) (Table I).

Spins were initialised uniformly inside (none outside) the simulated capillaries (5,000 per capillary); trajectories were generated with *δt* = 0.5 ms and *v* = 2 mm/s (giving *δx* = 1 µm).

Signals were generated using *δ* = 15 ms, Δ = 30 ms, 30 *b*-values uniformly spaced between *b* = 0 − 0.2 ms/µm^2^, and 32 gradient directions per *b*-value; signals were averaged over directions. The mono-exponential signal model (Eq. 11) was fitted to the simulated signals (averaged over the 10 phantoms in each set with the same morphology) using non-linear least squares to estimate *D*^*∗*^ values; 25 initial values were distributed uniformly within the constraints *D*^*∗*^ ∈ (0, 20) µm^2^/ms.

#### 2) Results

Fig. 4A shows the signals simulated in phantom sets with different segment lengths, *l*. Pseudo-diffusivity coefficients estimated from the simulated signals (Eq. 11) and the corresponding ground truth values (from Eq. 12) are given in Table I. There was excellent agreement between simulations and analytical solutions.

**Fig. 4.**
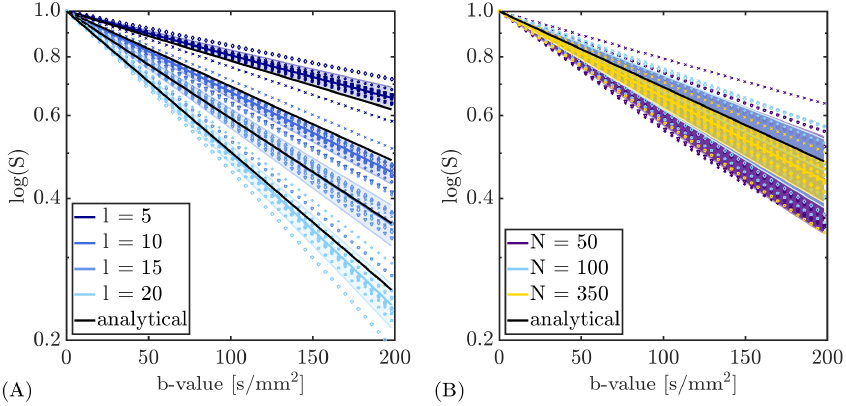
IVIM experiments: random-walk VASAL phantoms. Signal attenuation is shown for phantom sets with varying morphology. Markers represent individual phantoms, filled regions indicate the mean and SD over the set. Analytical solutions (Eq. 11) are in black. **(A)**. Phantoms with different segment lengths, ***l*** (***N* = 350** for all). **(B)**. Phantoms with different segment numbers, ***N*** (***l* = 10** ± **4** µm for all).

Fig. 4B shows signals simulated in phantoms with different segment numbers, *N* . Phantoms with fewer segments displayed higher variability in simulated signals and subsequently in the estimated apparent pseudo-diffusivities (Table I).

Note the spatial extent of continuous phantoms with *N* = 350 was *L* = 231 ± 32 µm^3^, indicating that *L* = 250 µm^3^ would be unlikely to introduce restriction effects in periodic phantoms and was appropriate for subsequent experiments.

### B. Non-Random Walks

#### 1) Methods

The effects of self-intersections, connecting angle statistics and vessel branching (i.e. deviations from phantoms generated as random walks) were investigated to assess the impact of network morphology on IVIM signals. Five periodic phantom sets (ten phantoms per set) - all with *N* = 350, *d* = 6 µm, *l* = 10 ± 4 µm, *L* = [250, 250, 250] µm^3^ - were created with the following morphological characteristics (Table I): phantom 1, self-intersecting with *g*_1_ = 1 and *θ*_*min*,1_ = 20^*°*^; phantom 2, self-avoiding, *g*_2_ = 1, *θ*_*min*,2_ = 0^*°*^; phantom 3, self-avoiding, *g*_3_ = 1, *θ*_*min*,3_ = 20^*°*^; phantom 4, self-intersecting, *g*_4_ = 5, *θ*_*min*,4_ = 0^*°*^; phantom 5, self-avoiding, *g*_5_ = 5, *θ*_*min*,5_ = 20^*°*^.

Spins were initialised uniformly inside (none outside) the simulated capillaries (5,000 per capillary); trajectories were generated using *δt*_*g*=1_ = 0.5 ms and *δt*_*g*=5_ = 0.2 ms for the *g* = 1 and *g* = 5 phantoms respectively (ensuring a maximum step size of *δx* = 1 µm), with *v*_*min*_ = 2 mm/s; flow velocities in the branching (*g* = 5) phantoms were therefore *v* = [2.0, 2.5, 3.2, 4.0, 5.0] mm/s (Eq. 14).

Signals were simulated as in Sec. VI-A.

#### 2) Results

Fig. 5 illustrates how each deviation from a random walk caused the signal attenuation and apparent *D*^*∗*^ to diverge from IVIM model predictions (Eq. 12); ground truth pseudo-diffusivities (from Eq. 12) and those estimated from the signal attenuation are given in Table I.

**Fig. 5.**
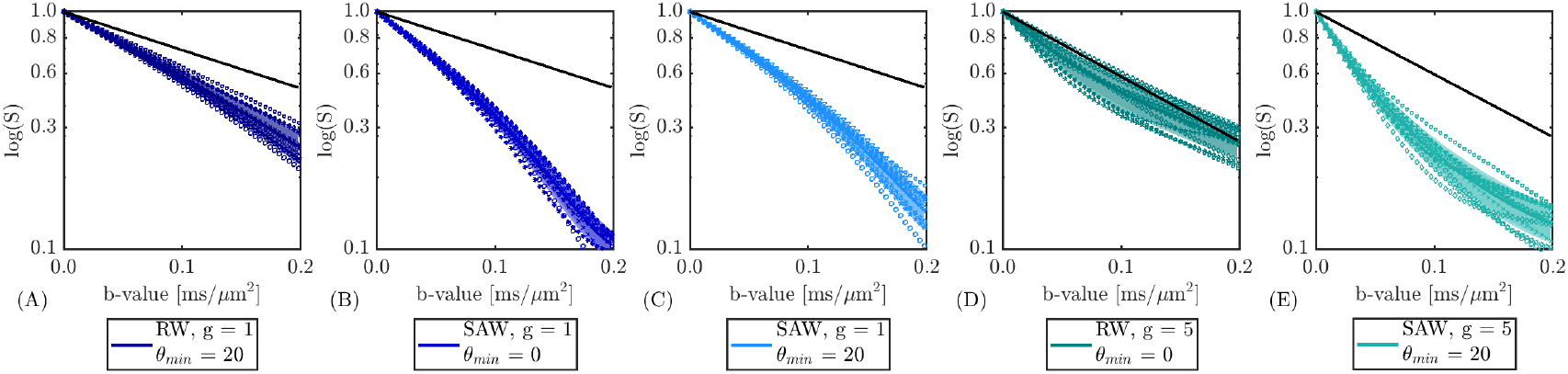
IVIM experiments: non-random walk VASAL phantoms. Signal attenuation plots are shown for the five phantom sets with different morphology. Markers represent individual phantoms; filled regions indicate the mean and SD over the set. Analytical solutions (Eq. 11) are in black. **(A)**. Self-intersecting (i.e. random-walk, RW) phantoms with ***g* = 1, *θ***_***min***_ **= 20. (B)**. Self-avoiding (i.e. self-avoiding walk, SAW) phantoms with ***g* = 1, *θ***_***min***_ **= 0. (C)**. SAW, ***g* = 1, *θ***_***min***_ **= 20. (D)**. RW, ***g* = 5, *θ***_***min***_ **= 20. (E)**. SAW, ***g* = 5, *θ***_***min***_ **= 20**.

Both self-avoiding phantoms and connecting angle constraints (i.e. *θ*_*min*_ *>* 0^*°*^) increased signal attenuation in simple phantoms without branching (*g* = 1) (Fig. 5A-C), with self-avoiding meshes additionally displaying non-Gaussianity at higher *b*-values (0.1 ≤*b* ≤0.2 ms/µm^2^).

The more complex phantoms with branching (*g* = 5) (Fig. 5D-E) also displayed non-Gaussian signal attenuation, even in self-intersecting meshes with *θ*_*min*_ = 0^*°*^ (Fig. 5D).

### C. Diffusion time dependence

#### 1) Methods

Diffusion time dependence was additionally explored in the phantoms comparable to the rat hippocampus (i.e. the periodic, self-avoiding phantoms with *g* = 5, *N* = 350, *d*_*min*_ = 6 µm, *l*_*min*_ = 10 ± 4 µm, *θ*_*min*_ = 20^*°*^, *L* = [250, 250, 250] µm^3^) (Table I).

Spins were initialised uniformly inside (none outside) the simulated capillaries (5,000 per capillary); trajectories were generated with *δt* = 0.2 ms and *v* = [2.0, 2.5, 3.2, 4.0, 5.0] mm/s across the generations (giving a maximum *δx* = 1 µm).

Signals were simulated with *δ* = 5 ms, Δ = [5, 10, 15, 20, 25, 30, 40, 50] ms, and two *b*-value ranges: (i) *b*_1_ = 0 − 0.05 ms/µm^2^, and; (ii). *b*_2_ = 0− 0.20 ms/µm^2^. For each range, 30 *b*-values were uniformly spaced between the bounds, with 32 gradient directions per *b*-value. Signals were averaged over all directions. The mono-exponential model (Eq. 11) was fitted (as previously described) to the simulated signals to estimate apparent *D*^*∗*^ values. The spherically-averaged DKI signal model, given by [35]:

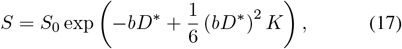

where *K* is the kurtosis, was also fitted, using 25 initial values uniformly distributed within the parameter constraints; these were *D*^*∗*^∈ (0, 20) µm^2^/ms, *K* ∈ (0, 3).

#### 2) Results

Fig. 6 shows the signal attenuation in VASAL phantoms for each diffusion time; the mean and SD of pseudo-diffusivity values (from the ADC and DKI models) and kurtosis (DKI model only) are given in Table II. Time dependence was observed at short times (Δ+*δ* ≲ 25 ms), where *D*^*∗*^ and *K* increased with time. Pseudo-diffusivities estimated using the DKI model were higher than from the ADC model; however, values were more comparable when using the lower *b*-value range (*b*_1_ = 0 − 0.05 ms/µm^2^) owing to increased kurtosis effects at higher *b*-values (*b*_2_ = 0 − 0.20 ms/µm^2^).

**TABLE II.**
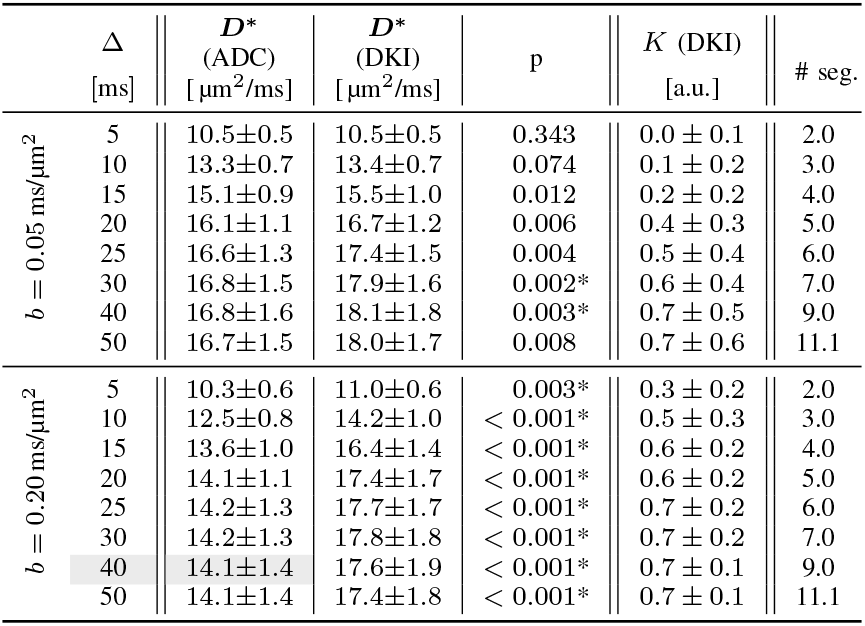
IVIM experiments: time dependence. Pseudo-diffusivities, ***D***^***∗***^, estimated using the ADC and DKI models are shown; significant differences (indicated by asterisks after the ***p***-values) were determined using a paired t-test with Bonferroni correction (***α* = 0.05*/*16 = 0.0031**). Kurtosis, ***K***, from the DKI model and the mean number of segments (# seg.) traversed during the experiment time are also shown.

**Fig. 6.**
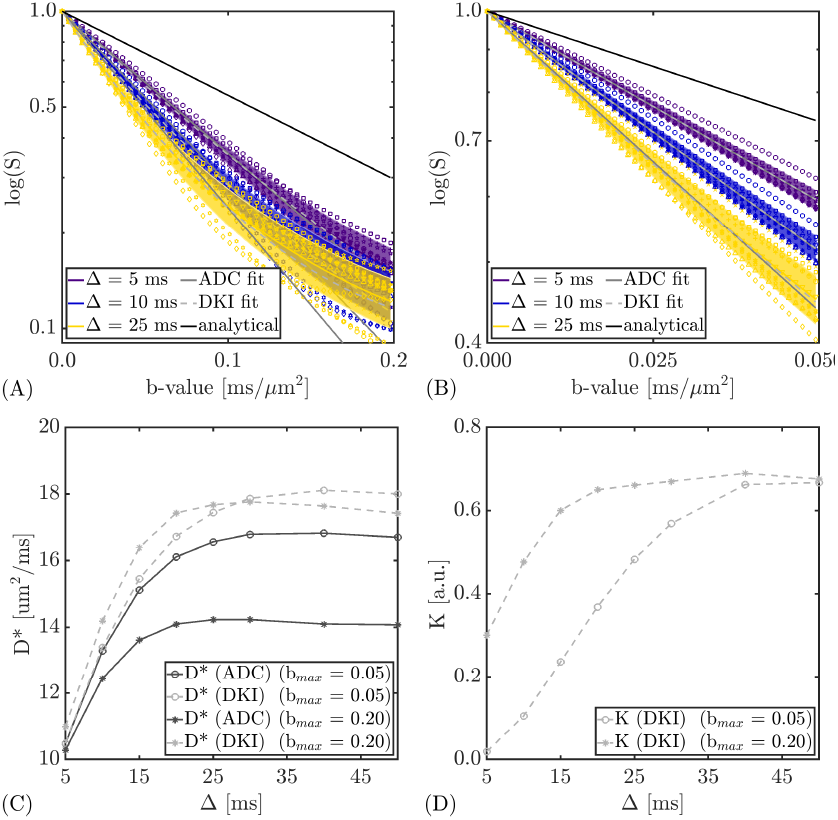
IVIM experiments: time dependence. **(A)**. Signal attenuation plots are shown for the VASAL phantom set resembling the rat hippocampus (periodic, self-avoiding, with ***g* = 5, *N* = 350, *d***_***min***_ **= 6** µm, ***l***_***min***_ **= 10** ± **4** µm, ***θ***_***min***_ **= 20**^***°***^, ***L* = [250, 250, 250]** µm^**3**^) at different diffusion times for ***b***_**2**_ **= 0** − **0.20** ms/µm^**2**^. Markers represent individual phantoms; filled regions indicate the mean and SD over the set. The ADC and DKI model fits are shown by the solid and dashed gray lines respectively; the analytical solution (Eq. 11) is in black. **(B)** As (A), but for the ***b***_**1**_ **= 0** − **0.05** ms/µm^**2**^ range. **(C)**. Pseudo-diffusivity values, ***D***^***∗***^, estimated using the ADC and DKI models for the two ***b***-value ranges are shown as a function of diffusion time. **(D)**. As (C), but for the kurtosis, ***K***, estimated using the DKI model.

## VII. Discussion

### A. A generative model for microvascular networks

Network morphology validation experiments demonstrated that VASAL phantoms replicating key features of rat hippocampal vasculature - such as branching patterns, segment lengths, diameters and orientations - could be generated (Fig. 2), highlighting the flexibility of VASAL. Generative models mimicking realistic microvascular geometries are an important step towards understanding the effects of microvascular alterations on MR signals; for example, changes in vessel branching, tortuosity, lengths and diameters, blood velocity, and perfusion fraction are indicated across a range of pathologies, including neurodegenerative disorders [36] and cancer [37], [38]. Using VASAL, it is possible to control each of these features independently, providing the potential to differentiate between vascular pathologies.

Perhaps the most comparable existing generative microvascular network algorithm is that of Linnegar et al. [31], which can synthesise the whole cortical circulation of the healthy mouse brain; however, as networks are grown using a global optimisation algorithm, simulating regional microvascular alterations (i.e. localised pathology) may be challenging. Moreover, while whole-cortex networks provide valuable multiscale information, their complexity may hinder analysis of how different microvascular geometries influence the MR signal. The smaller scale of VASAL phantoms can be utilised for this purpose, as demonstrated in Fig. 5.

A key feature of both algorithms, though, is their ability to generate closed phantoms with well-defined inlets/outlets, crucial for modelling blood flow and simulating dMRI signals. Existing IVIM simulation frameworks [23], [39] have so far used imaging-derived networks, which often lack clear boundaries. For example, the authors of SpinDoctor-IVIM [39] - a finite element modelling-based framework - chose a retinal network (derived from transmission electron imaging) for validation due to its limited number of inlets/outlets; however, spins required initialisation close to inlets to prevent their arrival at outlets during the simulation time, thereby limiting use of the entire network. SpinFlowSim [23] - an alternative framework using network pipe theory - used manually-segmented networks from histology and applied periodic boundary conditions by selecting inlets/outlets and creating shifted network replicas; however, the choice of inlet/outlet was random, and the networks were inherently 2D. Further, their small size (*N*∼ 55 segments) may lead to substantial variability in simulated dMRI signals. VASAL validation experiments indicated that smaller segment numbers, *N*, led to increased variability in simulated signals and subsequently increased uncertainty in estimated pseudo-diffusivities (Fig. 4B). Phantom segment numbers, *N*, must therefore be an important considerations when comparing networks with different morphologies.

### B. Observations from IVIM experiments

Using simple, true random-walk VASAL phantoms, synthesised MR signals were well approximated by the IVIM model (Fig. 4); however, deviations from perfect random-walk geometries caused simulated signals to diverge from the model predictions (Fig. 5), where phantoms with *θ*_*min*_ *>* 0 and SAW phantoms (even for *θ*_*min*_ = 0) all resulted in increased signal attenuation than predicted by IVIM. As both constraints prevent segments connected by smaller angles, local backtracking is minimised and causes an increase in the overall path length of the network, which subsequently increases the mean squared displacement of spins through the phantom and the corresponding signal attenuation. This suggests that, for realistic microvascular geometries, IVIM model assumptions may confound interpretation of measured *D*^*∗*^ values in terms of the underlying vascular microstructure, as altered geometries may also contribute to signal attenuation (and hence estimated *D*^*∗*^ values) in addition to segment length scales or blood velocities. It is unlikely that diffusion time dependencies contributed to these observations as spins were in the diffusive regime, traversing on average 9 segments in all phantoms over the experiment duration (Δ + *δ* = 45 ms) (Table I).

Diffusion time dependencies were observed, though, in VASAL phantoms at shorter times (Δ + *δ* ≲ 25 ms) (Fig. 6), where the increase in *D*^*∗*^ with diffusion time likely represented the increasing contribution of spins in the diffusive regime [40]. IVIM diffusion time effects have been previously reported in the brain [30], [40], with several alternative models proposed to account for spins in an intermediate regime between the diffusive and ballistic regimes [40]–[42]. Simulations here suggest that spins must travel on average more than 5 segments during the experiment to avoid diffusion time effects (Table II); however, this may also depend on phantom morphology and so requires further investigation in different geometries, which is beyond the scope of this work.

Kurtosis was observed across all diffusion times in simulated signals at higher *b*-values (*b* = 0.20 ms/µm^2^), and resulted in significantly greater *D*^*∗*^ estimates when using the DKI model compared to the ADC model (Table II). Kurtosis effects were less significant (high SD across phantom realisations) at lower *b*-values (*b* = 0.05 ms/µm^2^), although the DKI model still showed a tendency towards higher *D*^*∗*^ estimates. This suggests that the kurtosis observed in the bi-exponential decay of the IVIM dMRI signal around *b* = 0.1− 0.2 ms/µm^2^ - which is typically attributed to the intermediate regime between perfusion (high pseudo-diffusion) and tissue (low diffusion) effects - may also contain properties of the microvascular geometry. Disentangling these effects is likely to be challenging, but it is worth noting that signals at these *b*-values may contain as yet unexplored information on the microvascular network morphology.

### C. Potential applications

VASAL represents a unique tool to test the underlying assumptions of different acquisition and modelling paradigms. For example, the framework introduced here could be used to investigate, under different experimental conditions, which microvascular geometries can be detected or are negligible. There are also potential avenues using machine learning to disentangle different morphological changes: by simulating a large dictionary of different VASAL phantoms and corresponding MR signals, models could be trained to predict the underlying microvascular features from in vivo data. The VASAL phantoms mimicking the rat hippocampus here (Table I, grey highlight) took on average 4.5 hours to generate each; parallelised on 100 cluster nodes, then, it would take approximately 2-3 weeks to generate 10^4^ phantoms, which is a feasible timeframe for generating large training sets.

The Monte Carlo flow simulation framework could also be used for analysing real vascular examples, as it can accept as input any mesh with a pre-computed velocity field.

Lastly, although the focus here was dMRI, appropriate extensions of the Monte Carlo simulation framework may enable signal simulations for other techniques, for example dynamic contrast-enhanced MRI and arterial spin labelling.

### D. Limitations

Further validation of VASAL’s fidelity in replicating realistic microvascular morphology should involve simulating signals in real networks, such as the rat hippocampal example used here. However, substantive pre-processing is required for such complex networks; for example, inflow vessels must be identified or otherwise arbitrarily selected, velocity vector fields solved for the given inflow vessels, and physiologically accurate boundary conditions defined for all open vessel segments. Addressing these aspects was out of scope in this work, so validation was performed using geometric properties only; however, as it was demonstrated in simple, controlled networks that VASAL produced accurate dMRI signals (Fig. 3, 4), and assuming the rat hippocampal vessel morphology was accurately determined (Fig. 2), this validation was considered sufficient for exploring the theoretical aspects of IVIM here.

Simple plug flow was implemented in the Monte Carlo simulation framework presented here, neglecting any effects of laminar flow that may exist. As the aim of this work was to introduce and provide proof-of-concept of the novel VASAL tool, plug flow was deemed suitable for the experiments performed here, and indeed necessary under the SGP approximation, where diffraction patterns can be blurred by laminar flow. More complex velocity fields derived using fluid dynamics, for example, can easily be imported into the simulation framework in future studies, provided they are sampled sufficiently densely compared to the phantom size.

For similar reasons, extra-vascular spin dynamics and varying SNR levels were also not considered in the presented experiments; however, the toolbox allows simulation of both, which can be useful when, for example, investigating the ability of different modelling strategies to separate perfusion and diffusion effects.

Another limitation of VASAL is that the algorithm currently aims to generate segments with a random orientation distribution (as far as the user-defined angular constraints allow); however, vessels, particularly larger ones, may not always be randomly oriented, and may instead have a preferential orientation perpendicular to the sulcus surface [43], [44]. Adaptations to VASAL could be included to model such effects. Further, owing to the non-Markovian stochastic process of generating a SAW, creating large phantoms in small voxels may be challenging, as growing new segments without intersecting existing segments becomes increasingly difficult; a pragmatic solution may be to generate multiple smaller phantoms to achieve the desired perfusion fraction.

### E. Conclusions and future perspectives

A novel tool for generating microvascular phantoms – VASAL - and a framework for Monte Carlo flow simulations are introduced and demonstrated using select pseudo-diffusion examples. This work is a step towards realistic digital representations of brain microvasculature. Beyond addressing the limitations already discussed, a number of future developments are envisaged. For example, accurate exchange modelling is an important feature with potential implications for both IVIM and blood-brain barrier exchange imaging. Integrating VASAL phantoms with realistic cellular models [5], [7], [8] would also facilitate a more comprehensive representation of tissue microstructure.

The VASAL framework holds potential for advancing dMRI by enabling in-silico simulation of blood flow within physiologically accurate microvascular networks; this allows for rigorous testing and optimisation of novel dMRI sequences for sensitivity to specific microvascular features, for precisely interpreting the biophysical origins of subtle dMRI signal changes, for validating and refining biophysical models linking microvascular structure and function to dMRI biomarkers, and for investigating how pathological microvasculature alterations (e.g. in stroke, tumors, dementia, or small vessel disease) manifest in specific dMRI signatures for earlier and more specific diagnoses.

## Appendix A

### Geometric analysis framework for binary vessel data

The binary rat hippocampal micro-CT data were skeletonised (Matlab *bwskel*), then converted into a graph in which edges between nodes (voxels) were defined in a 26-voxel neighbourhood. The shortest paths between all terminal nodes - defined as voxels with degree = 1 or degree ≥ 3 - were computed using Dijkstra’s algorithm (Matlab *shortestpath*); these paths represent the curved vessel segment sets connecting all inlet/outlet pairs (degree = 1) or branch nodes (degree ≥ 3). Each curved segment was then discretised into a series of straight segments using a radius of curvature threshold of 0.25 (defined heuristically by visual inspection). The discretised segments were used to create a new graph, with nodes defined at each point between connecting straight segments (Fig. 7). This graph was used for all morphological feature analyses.

**Fig. 7.**
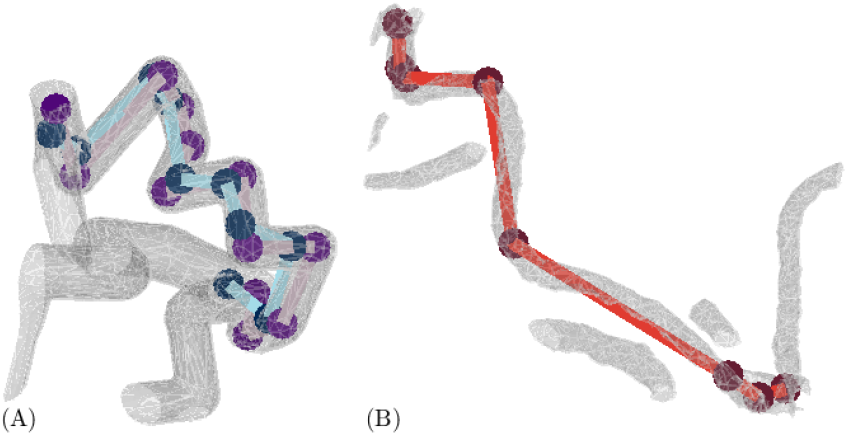
Estimating morphological characteristics. **(A)**. A small VASL phantom section is shown, with ground truth (purple) and estimated (blue) segment nodes and edges. **(B)**. A small section of the rat hippocampal mesh (created from the binary micro-CT segmentation) is shown, with the estimated segment nodes and edges (red).

Segment lengths were computed as the Euclidean distance between neighbouring nodes, and orientations as the azimuthal/elevation angles between segments and the *z*-axis. Segment diameters were calculated using ray intersections from the centre line (skeleton) with mesh faces (derived from the binary data using Matlab’s *isosurface*): the median distance to intersection across rays in a segment was taken as its radius.

## Acknowledgment

Thanks to Dr Laura Parkes and Dr Lauren Scott for providing the Wistar rat micro-CT data.

## Notes

E. Powell was supported by EPSRC grants EP/S031510/1 and EP/M020533/1. M. Palombo was supported by UKRI Future Leaders Fellowship grants MR/T020296/2 and 1073; MRC Research Grant MR/W031566/1; UKRI Research Grant BB/X005089/1.

### Competing Interest Statement

The authors have declared no competing interest.

